# A supervised learning algorithm to evaluate occurrence records in virtual species

**DOI:** 10.1101/2021.09.06.459158

**Authors:** Richard Rios, Elkin A. Noguera-Urbano, Jairo Espinosa, Jose Manuel Ochoa

**Affiliations:** Programa de Evaluación y Monitoreo, Instituto Alexander von Humboldt, Bogotá, Colombia; Universidad Nacional de Colombia, Sede Medellín, Facultad de Minas, Departamento de Energía Elećtrica y Automática, GAUNAL, Medellín, Colombia

**Keywords:** Point-occurrence records, expert knowledge, supervised learning, biological databases, virtual species, distribution modelling

## Abstract

Digital and open access of occurrence data have encouraged the development of tools to improve biodiversity conservation and management. In this study, we proposed a methodology to evaluate point-occurrence records based on expert knowledge. We firstly generated virtual data to test our methodology without confounding factors by simulating geographical distributions, virtual sampling, and expert checking of occurrence records. We used a set of non-linear bioclimatic variables and principal component analysis (PCA) to define a duality function between niche and biotope spaces. Subsequently, a supervised-learning model was fit to classify records between true and doubtful presence based on the virtual expert checking. We then tested our methodology using three virtual species and 10-fold cross validation. Also, we evaluated the prediction performance of the supervise model compared with the virtual observer using a virtual external database of occurrence data.

## Introduction

Occurrence data available in biological databases have expected errors that may reduce the quality of biographic studies [1–4]. Computational tools have been developed to assist the process of adding georeferencing information, as well as to detect and correct errors during the aggregation process [3, 6]. However, all the issues cannot automatically be detected and assistance of community experts is time consuming, tedious, or sometimes not available [2, 3]. In this setting, efforts should be focused in developing new tools and methodologies to improve the quality of occurrence data, as well as to integrate these tools into the best practice routines of data aggregation and management [2, 3].

In this study, we sought to develop a methodology to classify point-occurrence records as true and doubtful presences based on expert knowledge. Based on Hutchinson’s duality, we simulated virtual species using PCA with a set of non-collinear bioclimatic variables. Geographical distributions, virtual sampling, and expert checking of occurrence records of virtual species were simulated, seeking to test our methodology without confounding factors.

## Materials and methods

Our aim was to simulate realistic species distributions based on the Hutchinson’s duality [7]. Figure 1 provides the general simulation and analysis pathway used to develop the learning methodology. We used a set of non-linear bioclimatic variables and principal component analysis (PCA) to define a duality between niche and biotope spaces, where principal components served as the axes of the species’ niche. Base on the biotope-niche duality, we used the framework developed by Leroy et al. 2016 [8] to simulate virtual occurrence data, expert knowledge, and observation procedure. We build learning models to evaluate the suitability of occurrence records collected by a virtual observer based on the virtual expert knowledge.

**Fig 1.**
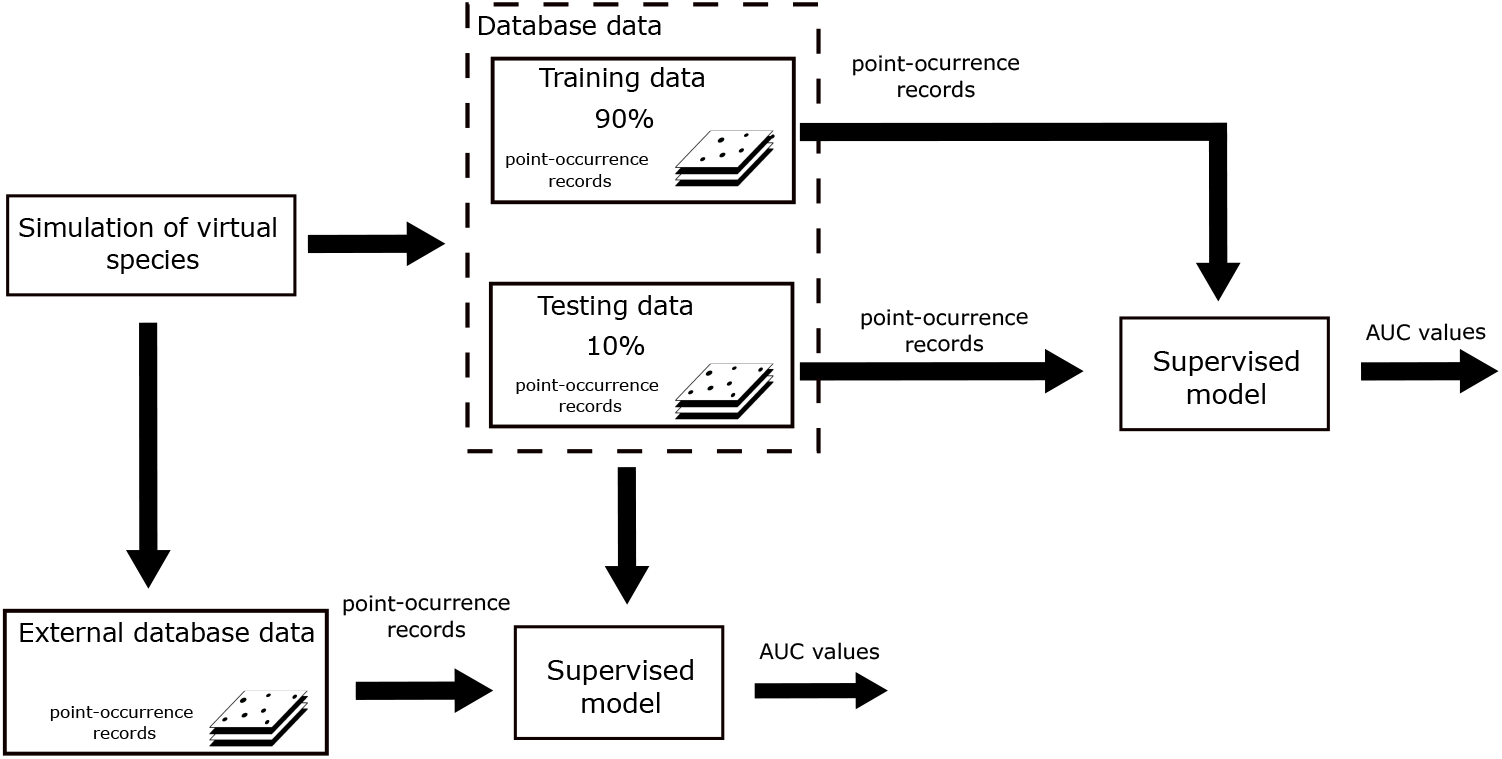
Supervised learning workflow to evaluate point-occurrence records in virtual species. Hutchinson’s duality and PCA was used to simulate virtual species with a set of non-collinear bioclimatic variables. Geographical distributions, virtual sampling, and expert checking of occurrence records of virtual species were simulated using “virtualspecies” package in R.

### Study area and data

Our study area was the Colombia’s continental territory. We used eight bioclimatic variables obtained from WorldClim at a spatial resolution of 1-km2, see Table 1 [9]. Bioclimatic variables were used to describe the Colombia’s bioclimatic space. The variable EnvOutlier described whether a bioclimatic variable had an outlier value: If a point record had an outlier value in at least one variable, then EnvOutlier was equal to 1; otherwise, the value of EnvOutlier was 0. We used only bioclimatic variables as climate is a major factor that describes distribution limits of several taxa and biota distribution [10, 11].

**Table 1.** Bioclimatic variables used in this study with their respective symbols.

| Bioclimatic variable | Symbol |
| --- | --- |
| Mean diurnal range | *Bio*<sub>2</sub> |
| Isothermality | *Bio*<sub>3</sub> |
| Temperature seasonality | *Bio*<sub>4</sub> |
| Min Temperature of coldest month | *Bio*<sub>6</sub> |
| Annual Precipitation | *Bio*<sub>12</sub> |
| Precipitation of wettest month | *Bio*<sub>13</sub> |
| Precipitation of driest month | *Bio*<sub>14</sub> |
| Longitude | Longitude |
| Latitude | Latitude |
| Altitude | Altitude |
| Environmental outlier | EnvOutlier |

### Simulation of virtual data

We simulated virtual data using a freely available package for R called “virtualspecies” to test our methodology without confounding factors related to intrinsic traits of real species, sampling process, and expert checking,see additional details in online tutorial at virtualspecies-tutorial. [8]. We simulated three virtual species with different types of environments. Below, we described the steps followed to generate the virtual data. Table 2 provides the parameters used to simulate occurrence data for each species.

**Table 2.** Virtual species’ simulation parameters. Virtual species were generated using the PCA based approach of the virtualspecies package implement in R.

| ID | Environmental suitability |  | Conversion into<br>absence and presence | Sampling<br>occurrence | Sampling area<br>delimitation |
| --- | --- | --- | --- | --- | --- |
|  | model 1 | model 2 | model 3 | model 4 | model 5 |
| 1 | 42/0.8 | -0.05/0.2 | 0.65/-0.05 | 0.9/0.2 | -73,-70.5,10.5,12.8 |
| 2 | -6.4/1.5 | 2.8/1 | 0.5/-0.05 | 0.9/0.2 | -78.8,-76,1,7 |
| 3 | -1.1/0.4 | -1.5/0.5 | 0.55/-0.05 | 0.9/0.2 | -76.8,-73.2,-4.2,8.3 |

### Virtual species distribution

We followed the three major steps proposed by Leroy et al 2016 to generate virtual species distributions using a PCA approach to ensure that each virtual species followed realistic environmental conditions. Firstly, we applied PCA to the set of bioclimatic variables to define a response function (i.e. the biotope-niche duality function) that described the species’ habitat suitability. Species-environment relationship was then simulated using Gaussian multivariate functions defined on the first two PCs, which served as the axes of the species’ niche space. Secondly, the environmental suitability was converted into a probability of occurrence based on a probabilistic approach with logistic functions. The probability of occurrence provided the probability of getting a presence of a species in a given background site [8]. Later, a random draw weighted by the probability of occurrence was performed to determine whether a background site was turned into a presence or absence. Thirdly, dispersal limitations were introduced to preclude the realized virtual distribution of each species into a subset of its potential distribution. An extent square area was defined to preclude any presence in background sites outside of this area.

### Virtual occurrence sampling

We simulated sampling occurrence data using the presence-absence scheme, which enabled to generate “real” and “observed” data. This scheme provides four variables: the point coordinates x and y, real presence*/*absence (1*/*0) of the species in the sampled background site (pixel) *real*, and the result of the sampling *observed* 1= presence, 0= absence. We considered the “real” variable as the labels provided by a virtual expert about true presences and absences of the species in the sampled background sites while “observed” variable was considered as observed presence and absence of a species resulting from historical, observational, or survey data [2].

This scheme enabled to introduce sampling biases, such as error and detection probability, to add errors between “real” and “observed” occurrence data. Differences between “real” and “observed” data can then be considered as commission (“real”=0 and “observed”=1) or omission errors (“real”=1 and “observed”=0), see more details in on-line tutorial [5, 8]. We then assigned a probability for detecting a generated virtual species in any background site weighted by the environmental suitability, i.e., less suitable background sites will have a lesser probability to detect the species. The probability of detection ranged from 0 (species could not be detected, i.e., “real”=0) to 1 (species could always be detected, i.e., “real”=1). Besides, we introduced a probability of having an erroneous detection of virtual species in a given background site. This scheme also enabled to restrict the sampling process to the spatial area defined to simulate the species’s geographical distribution, precluding the sampling occurrence points to the virtual species’s realized distribution. For each species, we randomly sampled two independent sets of point-occurrence data to validate the supervise model, hereby named as internal and external databases, respectively, i.e., we made two independent random draws of presence*/*absence points. The number of presence*/*absence points for each database was restricted to have 30% of presence records.

### Supervised model

We considered eight environmental and four geographical variables to build a supervised model that estimated the probability of an occurrence record of being a true presence, see Table 1. We used Extreme gradient boosting (XGBoost) to build the supervise model, which is leading technique in machine learning [12, 13]. XGBoost was implemented in R language (version 4.0.3) with the xgboost package (version 1.2.0.1). XGboost provides a rank of variable importance using the Gain metric, which describes the total loss reduction obtained for each feature during the tuning of model hyperparameters, i.e., it describes the relative contribution of each feature to the supervise model [12].

### Internal validation

Supervise model was initially trained and tested following an internal validation procedure with 10-fold cross validation. In the internal validation procedure, supervise model was tested 10 times with 10% of the data (test set) and trained with the remaining 90% of the data (training set). This procedure then resulted with 10 experimental models trained and tested with different sets, where only unseen data was used to test the supervise model. Validation results were concatenated to provide an estimate of the predictive performance when the supervise model could be applied to evaluate a new set of point-occurrence record. We used AUC to evaluate the prediction performance of the supervise model using the packages pROC (1.16.2) in R. PredictABEL (1.2.4) package was also used to generate the 10 random folds of the internal validation procedure.

### External validation

We also performed an external validation procedure to evaluate the supervised model, where internal database was used as training set while external database as test set. This validation procedure provided an estimate of the predictive performance when the supervise model could be applied to identify commissions errors in independent occurrence data [5]. We basically sought to evaluate the overall prediction performance of the supervise model to detect commission errors in occurrence data collected in field for a given species (“observed” data), which was previously labeled by an “expert” (“real” data). We excluded occurrence data where virtual species was detected (“real”=1) but not observed by the virtual observer (“observed”=0), seeking to exclude omission errors –sometimes produced when a species has with low detection probability. AUC and pairwise comparisons were used to evaluate the prediction performance of the supervise model and virtual observer to properly classify occurrence data [14]. Besides, we used the Youden Index to derive an optimal cutpoint for the supervise model to classify occurrence records as presences or absences for a given species [15]. We evaluated the improvement in specificity for the supervise model compared with the virtual observer –sensitivity analysis was omitted as omission errors were not considered for the virtual observer. Statistical analysis were performed using the pROC (1.16.2) and cutpointr (1.0.2) packages implemented in R.

## Results

### Simulation of virtual species

For species 1, species-environment relationship and environmental suitability are provided in Figure 2 and 3, respectively. In addition, map of probability of occurrence and occupancy for species 1 are provided in Figure 4 and 5, respectively. Similarly, occurrence data virtually sampled for species 1 is depicted in Figure 6.

**Fig 2.**
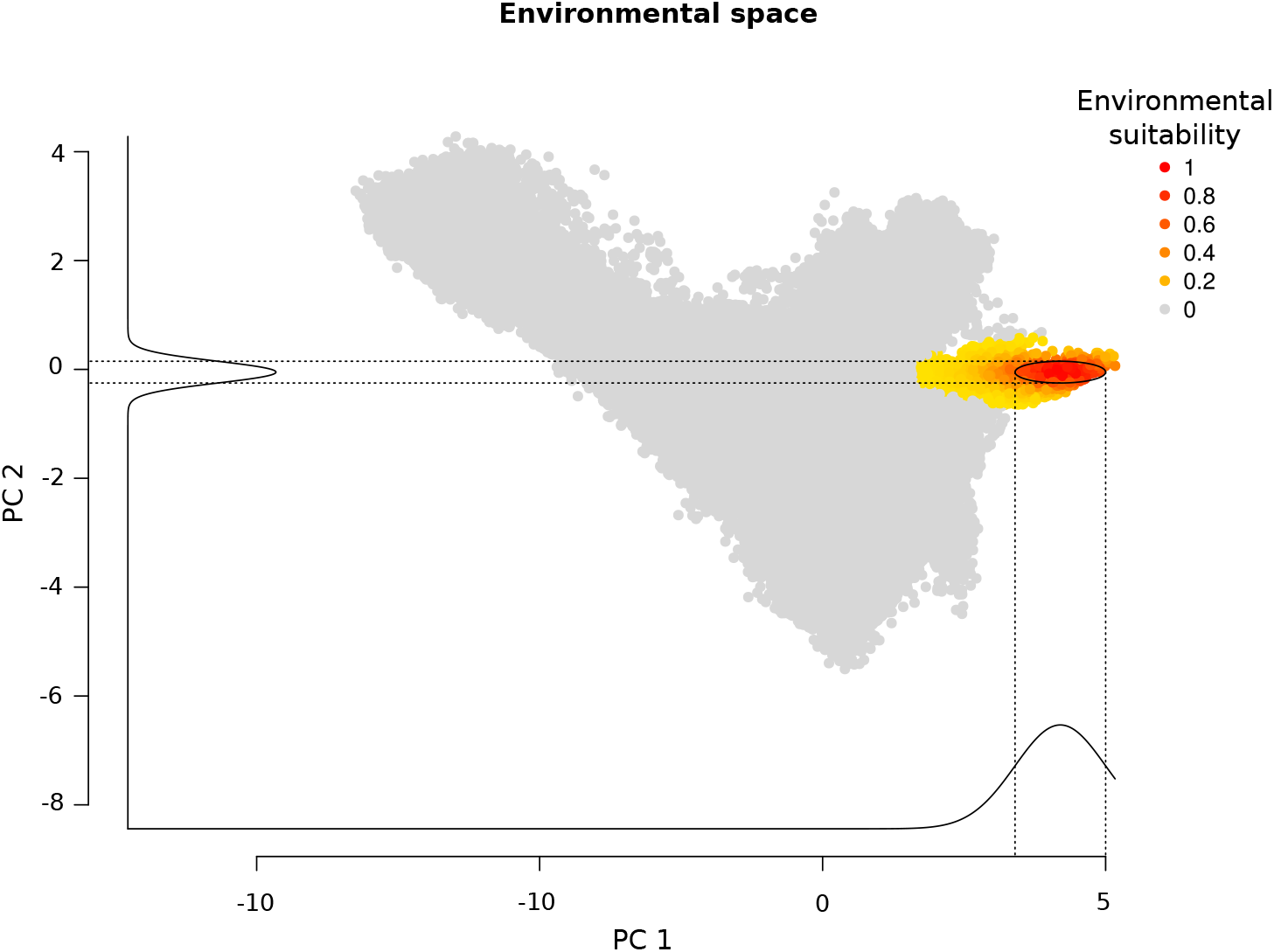
Species-environment relationship for species 1. Graphical representation of the species 1’s environmental conditions defined with Gaussian multivariate functions defined on the first two PCs.

**Fig 3.**
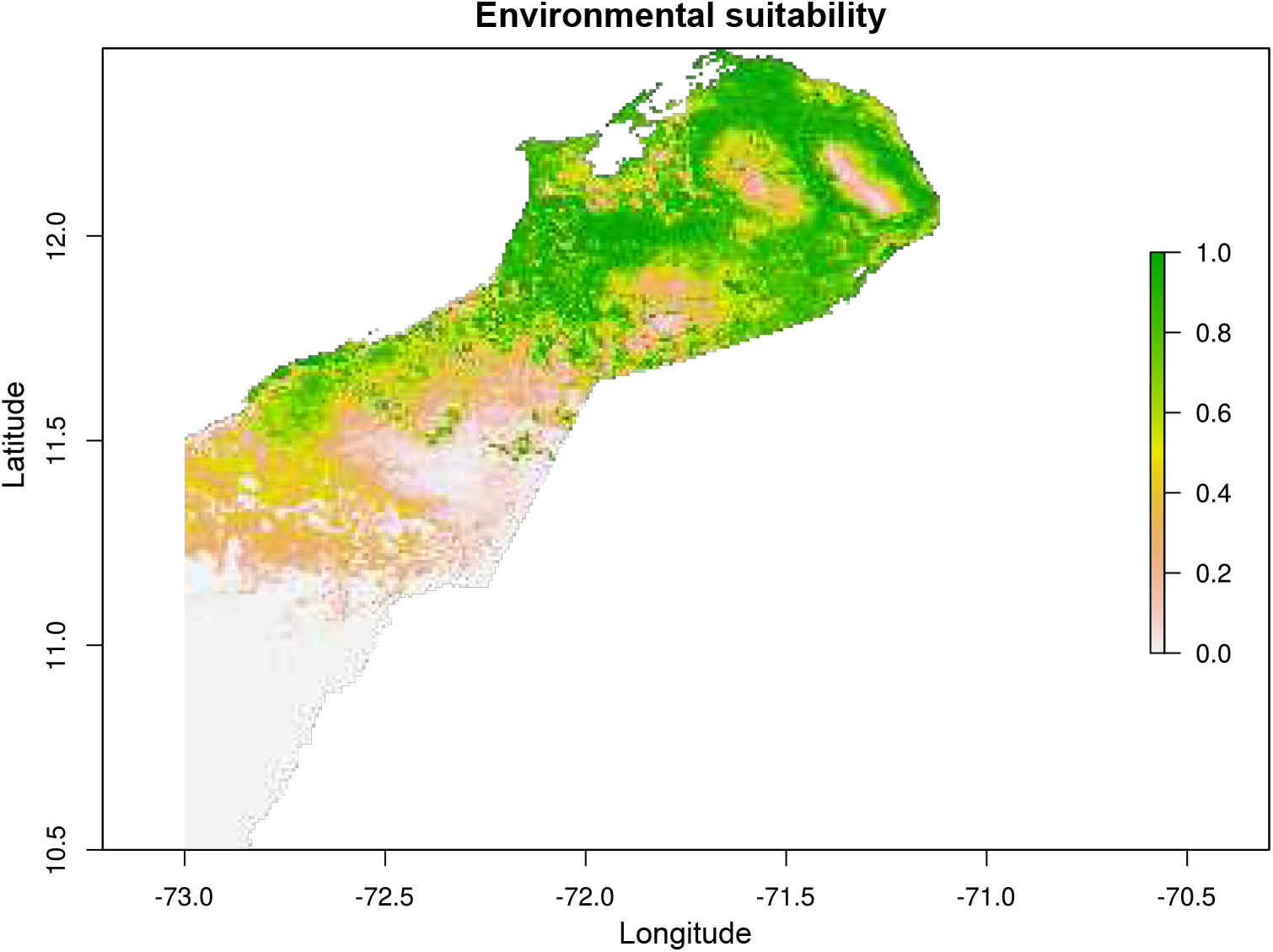
Environmental suitability for species 1. Predicted environmental suitability for the species 1 in the physical space restricted to its extent area.

**Fig 4.**
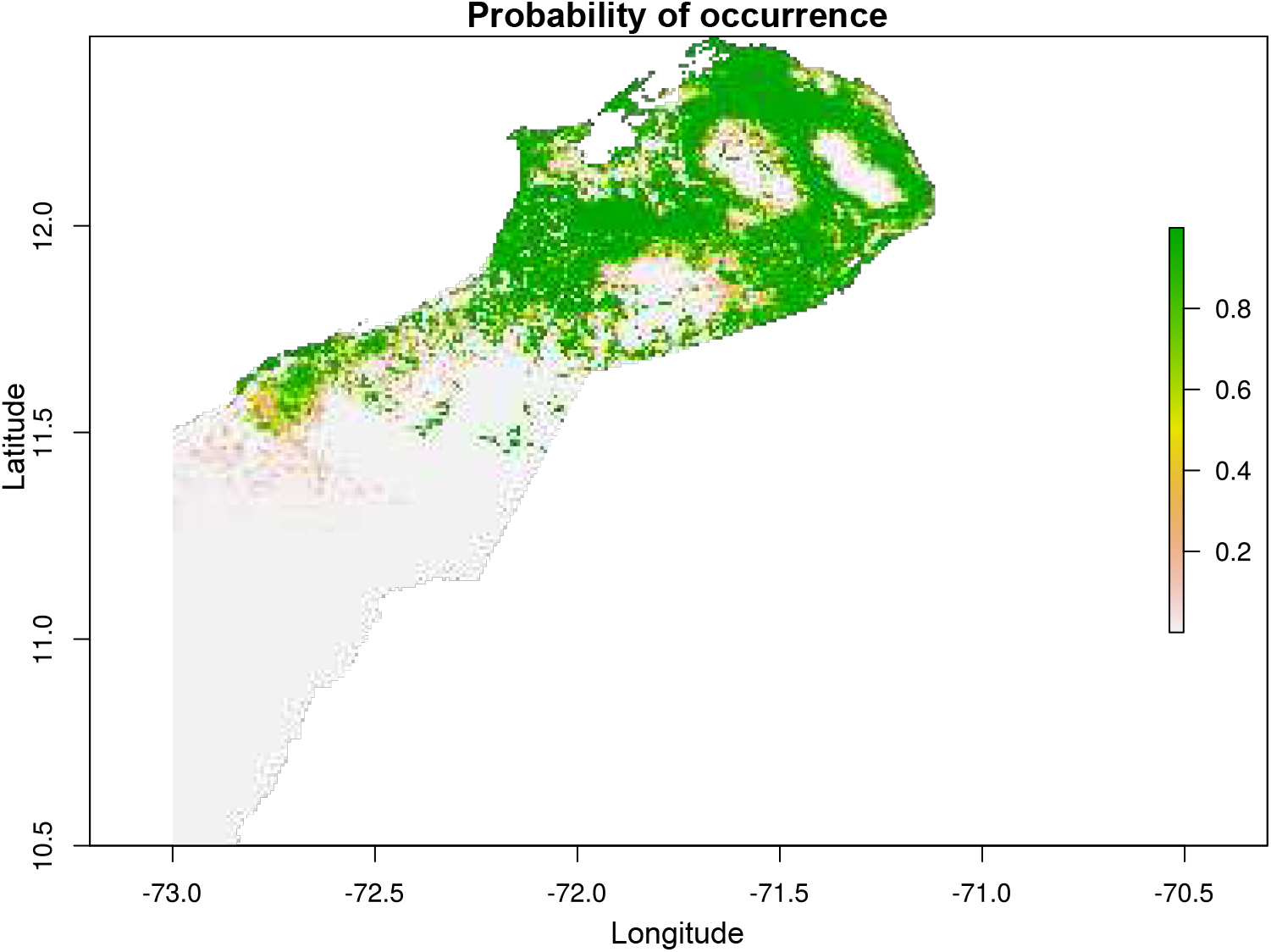
Map of probability occurrence for species 1. Each pixel described the probability of getting a presence of the species 1. This probability is estimated using a logistic function with the environmental suitability as independent variable.

**Fig 5.**
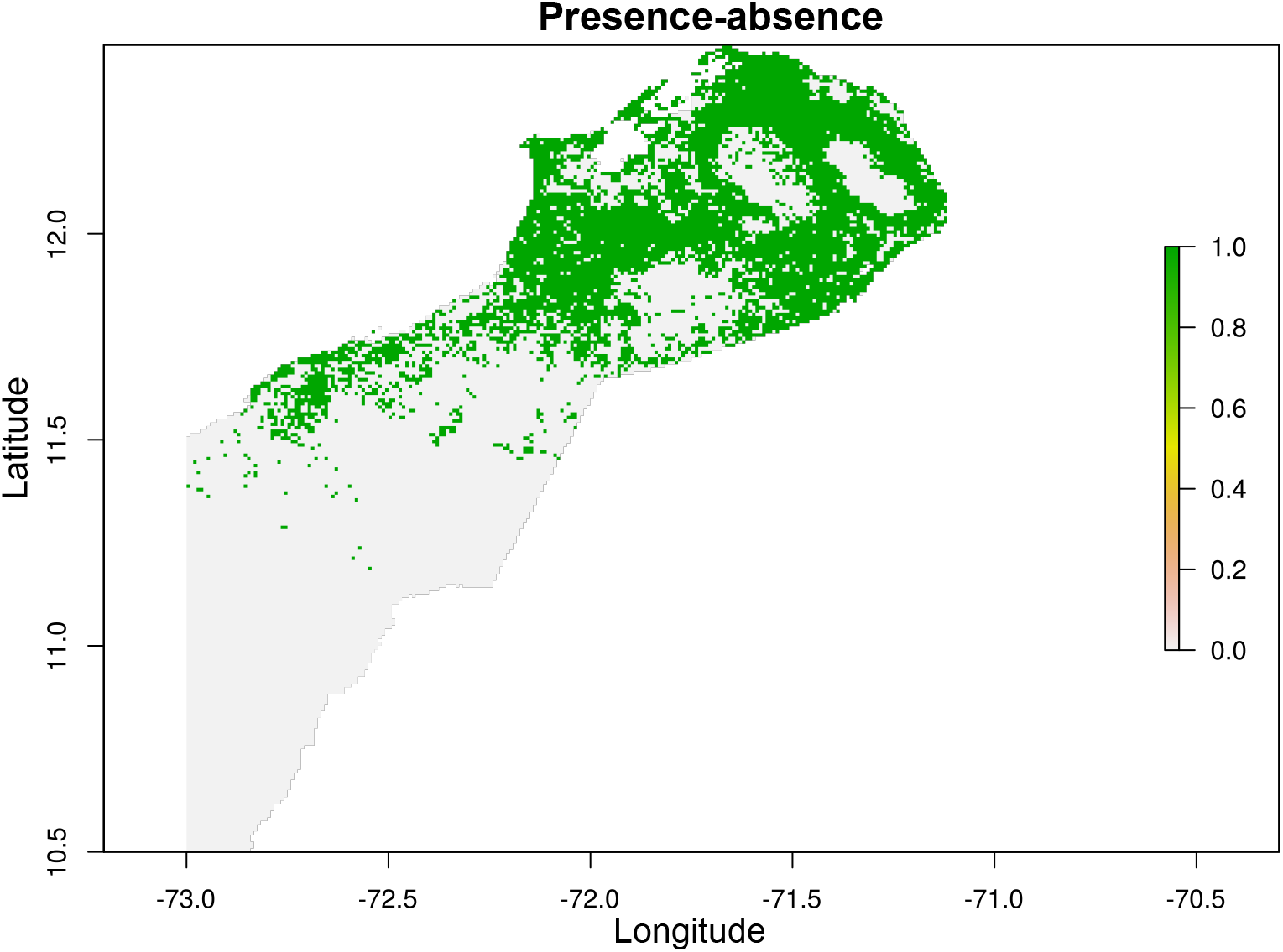
Occupancy map estimated for species 1. Realized distribution estimated for species 1 by performing a random sample of presence and absence weighted by the probability of occurrence.

**Fig 6.**
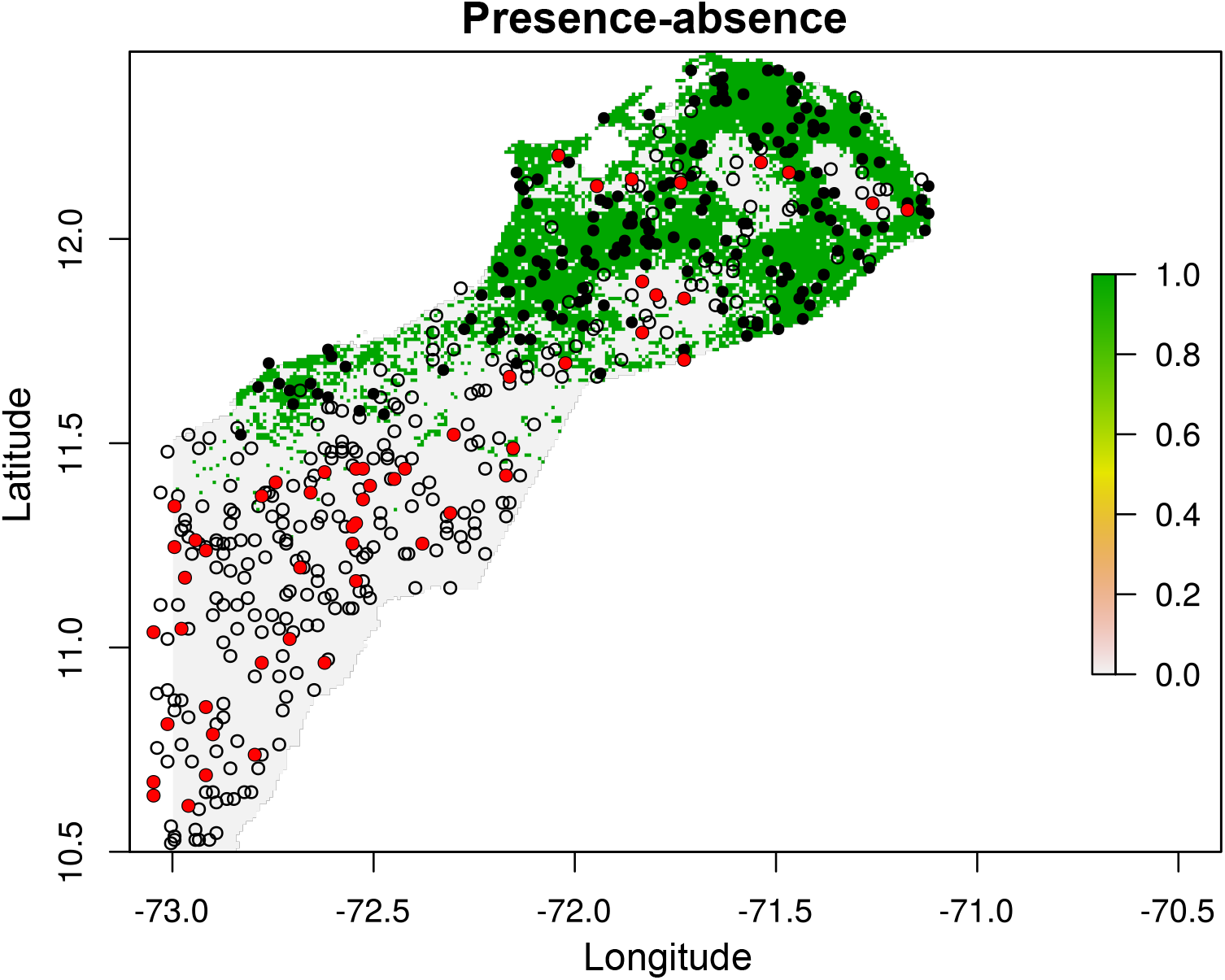
Occurrence data sampled for species 1 in the internal database. Green pixels present a realized distribution map estimated for species 1. Black points and empty circles represent background sites where the species was detected (“real”=1) and absent (“real”=0), respectively. Meanwhile, red points represent background sites where the species was present (“Observed”=1), but not detected (“Real”=0). Red points were considered as commission errors, either by inaccurate recording of geographic co-ordinates or taxonomic errors.

### Internal validation

Figure 7 provides the average importance ranking for all variables for species 1, which derived by averaging the ten rankings obtained from each training set during the internal cross validation procedure. The prediction performance of the species 1’s supervise model was AUC = 0.945, 95% CI: 0.927 to 0.964. Similarly, for species 2 and 3, the prediction performance of the supervise model was AUC = 0.995, 95% CI: 1 to 0.989 and AUC = 0.892, 95% CI: 0.948 to 0.836, respectively.

**Fig 7.**
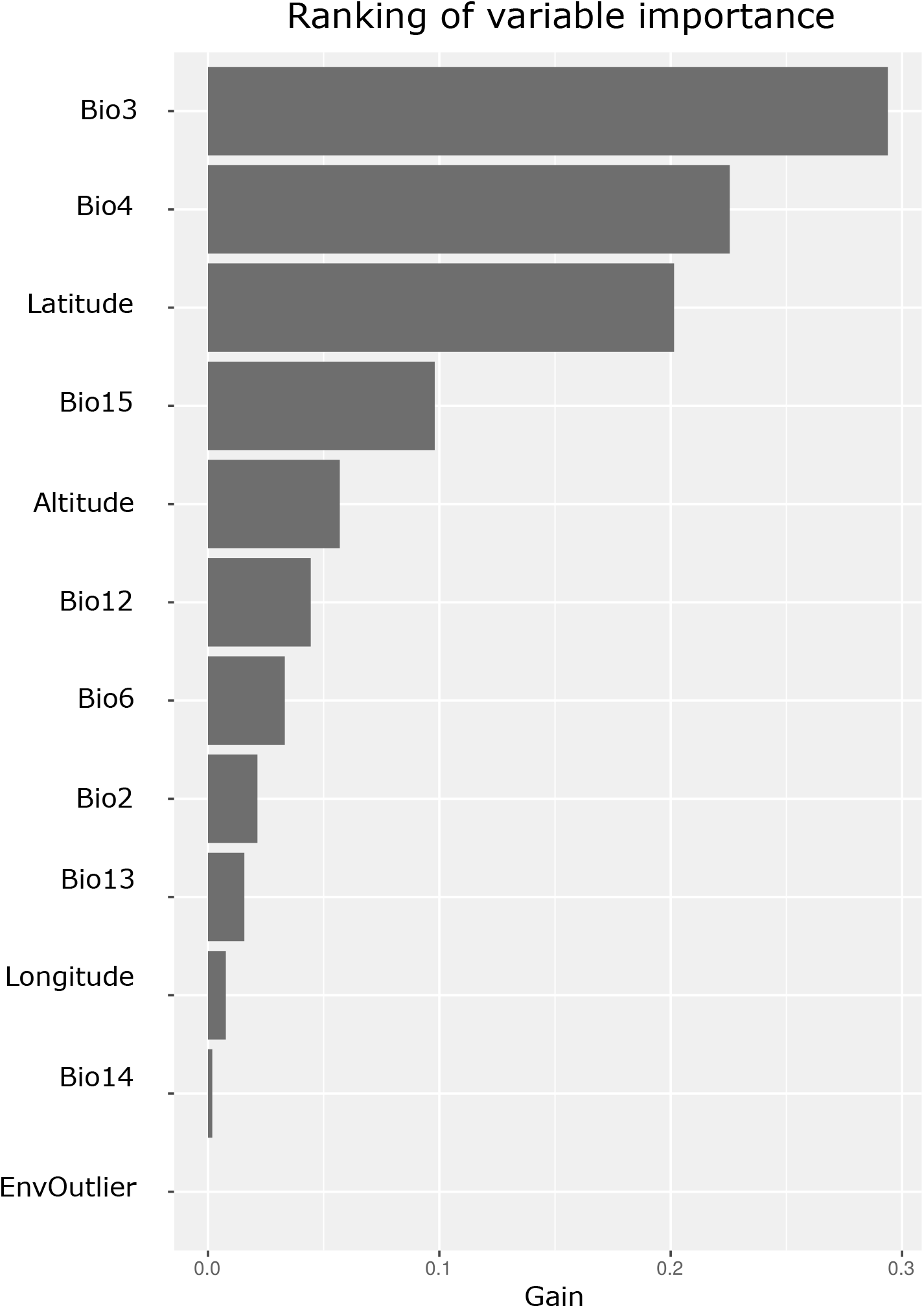
Ranking of variable importance for species 1. XGBoost’s gain metric was used to determine the contribution of each variable to the model. Importance ranking of all variables were derived by averaging the 10 rankings obtained each training set in the internal cross-validation procedure.

### External validation

We evaluated the prediction performance of the supervise model for detecting commissions errors in the external databases. Table 3 provides the prediction performance of the supervise model and virtual observer. When operating with Youden cutpoint, Table 4 provides specificity and sensitivity values for the supervise model compared with the virtual observer for each species. The optimal cutpoints were 0.532, 0.398, and 0.374 for the species 1, species 2, and species 3, respectively. Figure 8 depicts the occurrence data of the external database in the geographical space, as well as the true absences classified as true presences, either by the supervise model or virtual observer (i.e., the false positives). Similarly, Figure 9 depicts the occurrence data of the external database projected in the species 1’s niche space, as well as the false positives, either by the supervise model or virtual observer.

**Table 3.** External validation of the supervised model. AUC values for the supervise model and virtual observer in the external validation database. Abbreviations: AUC – area under the receiver-operating characteristic curve; CI –confidence interval.

|  | Virtual species 1<br>AUC (CI:L95-U95) | Virtual species 2<br>AUC (CI:L95-U95) | Virtual species 3<br>AUC (CI:L95-U95) |
| --- | --- | --- | --- |
| Observer | 0.896 (0.872-0.92) | 0.915 (0.884-0.945) | 0.898 (0.857-0.939) |
| Supervise model | 0.956 (0.939-0.974) | 0.994 (0.986-1) | 0.906 (0.852-0.961) |
| p-value | < 0.001 | < 0.001 | < 0.001 |

**Table 4.** Specificity analysis of the supervised model in the external database. AUC values for the supervise model and virtual observer in the external validation database. Abbreviations AUC – area under the receiver-operating characteristic curve; FN –false negatives; FP –false positives.

|  | Virtual species 1 |  | Virtual species 2 |  | Virtual species 3 |  |
| --- | --- | --- | --- | --- | --- | --- |
|  | Specificity /FP | Sensitivity /FN | Specificity /FP | Sensitivity /FN | Specificity /FP | Sensitivity /FN |
| Observer | 0.79/58 | 1/0 | 0.83/25 | 1/0 | 0.796/19 | 1/0 |
| Supervise model | 0.917/23 | 0.879/14 | 0.966/5 | 0.981/1 | 0.903/9 | 0.8/7 |

**Fig 8.**
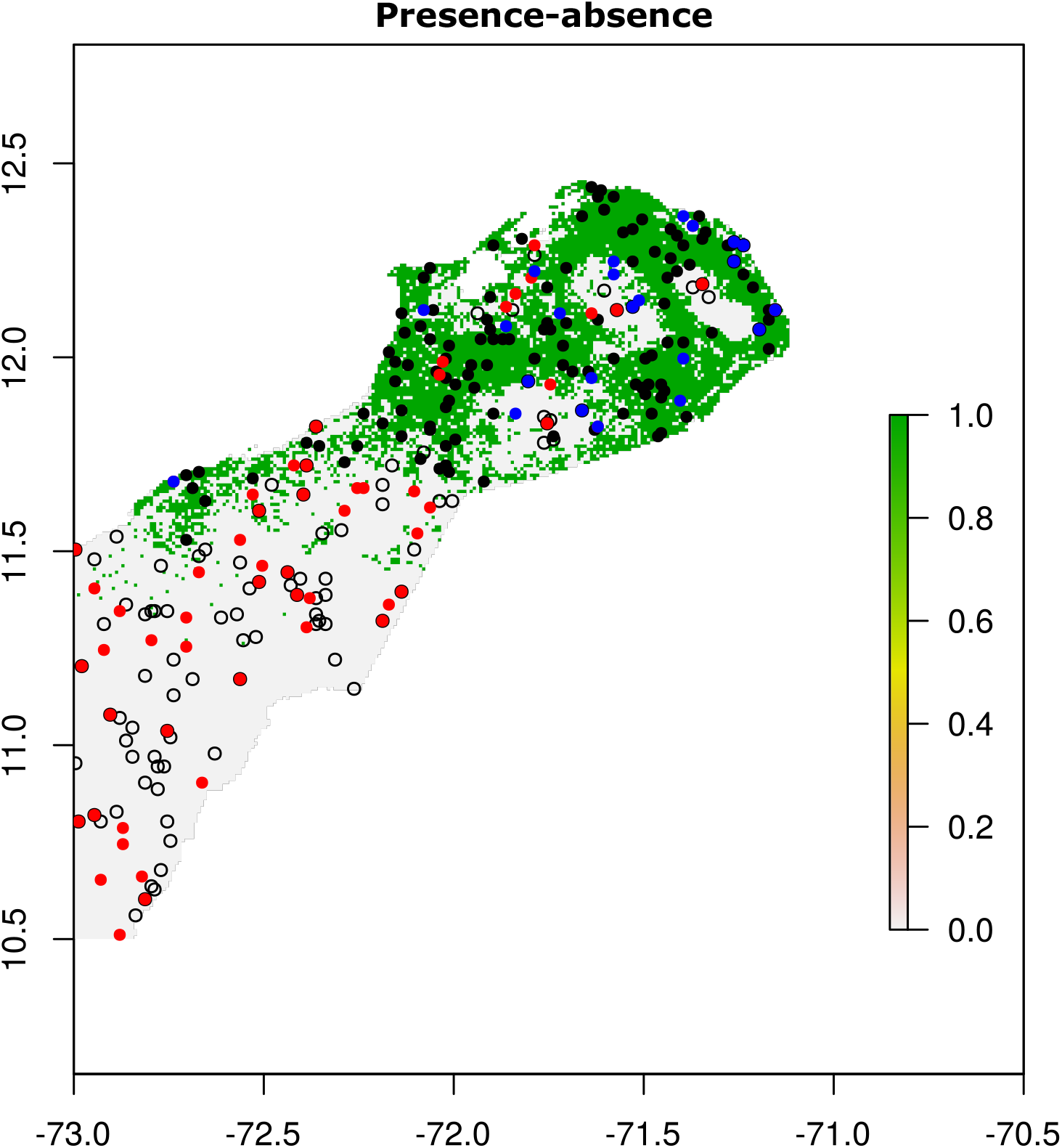
Occurrence data sampled for species 1 in the external database. Green pixels present the realized distribution map estimated for species 1. Black points and empty circles represent background sites where the species 1 was detected (“real”=1) and absent (“real”=0), respectively. Meanwhile, red points represent background sites where the species was present (“Observed”=1), but not detected (“Real”=0), i.e., red points represented commission errors made by the virtual observer, either by inaccurate recording of geographic co-ordinates or taxonomic errors. Similarly, blue points represent background sites where the species was predicted to be present by the supervise model but not detected (“Real”=0) with a cutpoint of 0.532, i.e., blue points represented commission errors made by the model.

**Fig 9.**
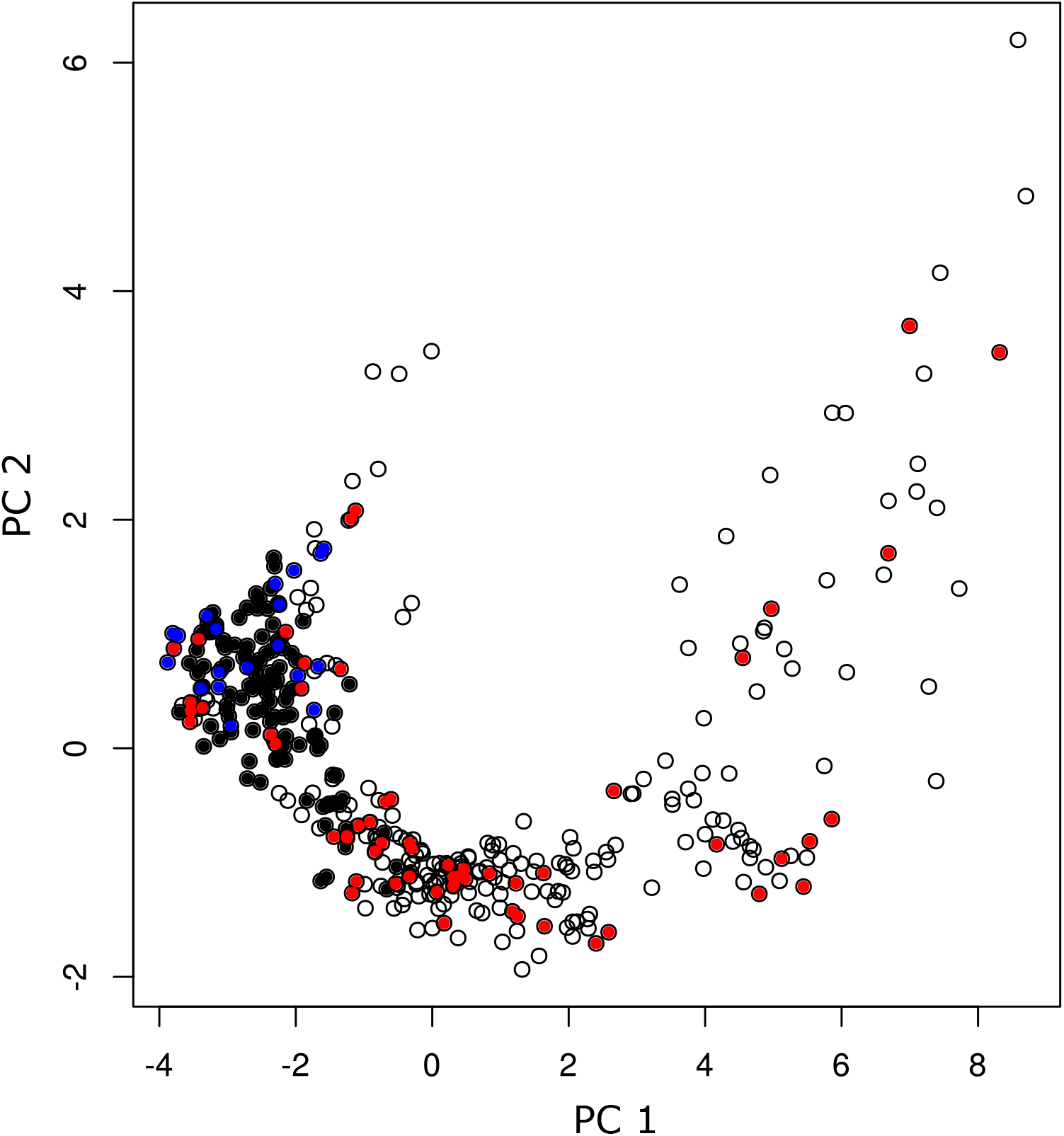
Eternal occurrence data of the species 1 projected in its niche space. Black points and empty circles represent occurrence data projected in the species 1’s niche space, where the species 1 was detected (“real”=1) and absent (“real”=0), respectively, according to occurrence data in the external database. Meanwhile, red points represent background sites where the species was present (“Observed”=1), but not detected (“Real”=0), i.e., red points represented commission errors made by the virtual observer, either by inaccurate recording of geographic co-ordinates or taxonomic errors. Similarly, blue points represent background sites where the species was predicted to be present by the supervise model but not detected (“Real”=0) with a cutpoint of 0.532, i.e., blue points represented commission errors made by the model.

## Discussion

In this study we proposed and evaluated a methodology to classify point-occurrence records between true and doubtful presences based on expert knowledge. We simulated virtual species to test our methodology without confounding factors based on Hutchinson’s duality, generating geographical distributions, virtual sampling, and expert checking of occurrence records. We showed that machine learning can approximate the niche space for a given specie by learning from occurrence data labeled by an (virtual) expert. We showed that machine learning models can reproduce expert knowledge and help to identify errors in occurrence data. ML models could be integrated into aggregation tools to automatically detect and correct errors in occurrence data [3].

## Conclusion

In this study we have proposed a methodology to predict true presences in point-occurrence records based on expert knowledge. We sought with this methodology to learn from expert opinion to develop a computational tool that automatically evaluates occurrence records and identifies commission errors. These tools can be integrated into recommendation systems to assist the aggregation process of georeferencing information into biological databases.

## Acknowledgement

This research was supported by the Departamento Administrativo de Ciencia, Tecnología e Innovación (COLCIENCIAS) – Colombia with the “ Programa de Estancias Postdoctorales Beneficiarios COLCIENCIAS 2017 ” grant.

## Notes

### Competing Interest Statement

The authors have declared no competing interest.

